# Uptake of environmental DNA in *Bacillus subtilis* occurs all over the cell surface through a dynamic pilus structure

**DOI:** 10.1101/2023.03.10.532016

**Authors:** Alexandra Kilb, Marie Burghard-Schrod, Sven Holtrup, Peter L. Graumann

## Abstract

At the transition to stationary phase, a subpopulation of *Bacillus subtilis* cells can enter the developmental state of competence, where DNA is taken up through the cell envelope, is processed to single stranded DNA, which is incorporated into the genome if sufficient homology between sequences exists. We show here that the initial step of transport across the cell wall occurs via a true pilus structure, with an average length of about 500 nm, which assembles at various places on the cell surface. Once assembled, the pilus remains at one position and can be retracted in a time frame of seconds. The major pilin, ComGC, was studied at a single molecule level in live cells. ComGC was found in two distinct populations, one that would correspond to ComGC freely diffusing throughout the cell membrane, and one that is relatively stationary, likely reflecting pilus-incorporated molecules. The ratio of 65% diffusing and 35% stationary ComGC changed towards more stationary molecules upon addition of external DNA, while the number of pili in the population did not strongly increase. These findings suggest that the pilus assembles stochastically, but engages more pilin monomers from the membrane fraction in the presence of transport substrate. Our data reveal that localized transport of DNA into the cytosol at a single cell pole is preceded by localization throughout the periplasm and efficient uptake through the entire cell surface by a dynamic pilus.

**Importance:** Horizontal gene transfer (HGT) is important for bacterial evolution and the adaption to new environments. Through HGT, genes can be transferred between bacteria, which can lead to antibiotic resistance development, especially critical with regards to pathogenic bacterial species. One mechanism of HGT is natural competence, through which bacteria can take up DNA under natural conditions and incorporate it into their chromosomes. For DNA uptake, a complex will be formed that is conserved among Gram-positive and Gram-negative bacteria, in spite of different envelope architectures. We show that in *Bacillus subtilis*, where the part of the uptake complex spanning the cell membrane localizes to a single cell pole, transport through the cell wall is mediated by a visible pilus structure that can extend and retract to pull DNA into the periplasm. Thus, DNA uptake from the environment is not limited to polar sites, but is channelled into the cytosol at the pole. Single molecule dynamics of ComGC revealed a static, likely pilus-bound, and a mobile fraction of ComGC within the cell membrane. Addition of environmental DNA increased the static fraction of molecules, showing that pilus dynamics respond to DNA binding.

## Introduction

Prokaryotes are known to colonize almost every place on planet earth, including environments with extreme environmental conditions that require specific traits to survive [1]. Bacteria present the largest domain of life, and are known for their adaptability to changing environmental conditions. There are two main reasons for genomic adaptation; one is horizontal gene transfer (HGT), the other one is accumulation of beneficial mutations based on random base changes in DNA, within large population sizes. HGT is facilitated by the uptake of foreign DNA, by conjugation (direct transfer involving a pilus structure), transduction (via phage infection) or competence [2]. Naturally competent bacteria can take up environmental DNA through the cell envelope, and become transformed when DNA is incorporated into their genome, which is a physiological and genetical trait of many bacterial strains [3]. Alternatively to chromosome integration, taken up DNA can be recombined to yield plasmid DNA, in case such DNA has been available [4], or can be metabolized in the cell as a source of phosphate, nitrogen and carbon [2, 5, 6]. Some bacterial strains, including pathogens, are naturally competent, and can thus obtain antibiotic resistance genes through natural transformation leading to drug resistance; therefore, competence is of clinical relevance [7, 8]. In 2014 it was reported that 80 bacterial strains were identified to be naturally competent under various conditions, but likely, there are many more species that can become competent under conditions that are not yet known [9]. The Gram negative bacterium *Vibrio cholerae* takes up exogenous DNA on chitin surfaces [10, 11], while *Neisseria gonorrhoea* shows constitutive expression of competence [12]. In other bacteria, like in the Gram-positive *B. subtilis* or the Gram-negative bacterium *Haemophilus influenzae*, the development of competence is induced by e.g. conditions that limit growth (reviewed in [3, 13]). This variability of competence-induction has the purpose to serve the special needs for the organisms [3]. In some organisms, like the human pathogens *N. gonorrhoeae* and *H. influenzae*, preferences for specific DNA uptake sequences were identified, which in this case consist of 10 base pairs [14]. In *B. subtilis*, sequence preference for uptake is not known. During the state of competence around 100 genes are upregulated in this organism, of which 20 are essential for DNA uptake [15, 16]. Environmental DNA is taken up by a multiprotein complex, which differs between bacterial strains, but is similar in its basic structure. In gram-negative bacteria, DNA has to overcome an additional outer membrane, while in Gram positive bacteria DNA has to pass a thick 30-50 nm peptidoglycan layer [17]. The proteins forming the uptake complex in *B. subtilis* were identified in a systematic transposon screen [18]. It is formed in just 2-20% of cells within a culture, dependent on the strain [19]. Thus, cell fates such as competence or sporulation are active decisions, which are pursued by a subpopulation of cells. Decisions are based on bistable switches, an intriguing phenomenon in bacterial populations [20]. The regulatory system of competence is induced via quorum sensing, i.e. by secreted peptide factors produced by the cells, leading to the stabilization of the master transcription factor ComK, whose gene, *comK*, is repressed by *rok* (<u>R</u>epressor <u>o</u>f <u>c</u>ompetence). The deletion of *rok* results in up to 80% of cells reaching the state of competence [21]. The DNA uptake complex is composed of late competence proteins encoded by four different operons [22–25]. Intriguingly, parts of the machinery have been shown to be located at one, or rarely at both cell poles [26], showing that DNA uptake – at least through the cell membrane - occurs at a specific site in competent cells. Accumulating evidence suggests that Com proteins localize specifically near the poles due a diffusion and capture mechanism [27, 28], with a true “anchor” protein still being unknown. When cells are converted into round spheroplasts, single positions within the membrane were found to persist, revealing that the assembly is highly stable and does not require the cell pole for its maintenance [29]. Immunofluorescence microscopy revealed that, like other competence proteins, ComGC, the major pilin forming the basis of the pseudopilus structure, also localizes at poles and cell centre. Therefore, it was proposed that cells may contain just one active, assembled competence machinery and non-assembled single competence proteins [29].

Common to DNA uptake systems in Gram negative bacteria is that the periplasmatic uptake seems to take place by retraction of a pilus structure [2]. For *B. subtilis*, the existence of a “pseudopilus” was shown, composed of a major pilin (ComGC) and some minor pilins (ComGD, ComGE, ComGG). These membrane proteins are processed in their N-terminal region to gain the ability of formation of the pilus structure, by ComC, the pre-pilin peptidase. An intramolecular disulphide bond is introduced into ComGC by an oxidoreductase pair BbbD und BdbC [30]. Loss of the cell wall relieves the essential nature of the ComG operon for DNA uptake [22], suggesting that the complex, whose nature is still unclear, is essential to move DNA through the cell wall. In other bacteria, pili have been labelled *in vivo* by introducing a cysteine substitution in the major pilin and subsequent labelling with maleimide dyes [31]. With this method, pilus structures in the human pathogens *V. cholerae* or *Streptococcus pneumoniae* were identified and stained. It was possible to follow the DNA uptake in real time and it was shown that pilus retraction is essential for DNA uptake in these bacteria [32, 33]. Contrary to gram-negative systems, a retraction ATPase is missing within the systems of gram-positive bacteria. It was therefore proposed that also ComGA, the assembly ATPase in *B. subtilis*, might have a bifunctional role, or that cytosolic DNA helicase ComFA might induce retraction, facilitating the uptake of environmental DNA [2]. After uptake into the periplasm, double stranded DNA binds to the DNA receptor ComEA at its C-terminus [34, 35]. This domain is composed of Helix-Hairpin-Helix motives which interact with DNA without any sequence specificity. In recent work, tracking of fluorescently labelled DNA in *B. subtilis* showed that taken up DNA is localized in the periplasm before it enters to the cytosol [28, 36]; the periplasm thus acts as a reservoir for DNA like in *V. cholerae* [37]. The DNA might itself be mobile within the periplasm or might move together with ComEA. After degradation of one DNA strand, single-stranded DNA can enter to the cytosol through the channel protein ComEC, which is localized at the cell pole in competent cells, and may have nuclease activity *in vivo* for generating ssDNA [38]. A key missing question is the localization and architecture of the complex mediating cell wall passage of DNA in *B. subtilis*, which we show to have true pilus properties and to respond to interaction with environmental DNA.

## Methods

### Growth conditions

*Escherichia coli* and *Bacillus subtilis* cells were grown in lysogeny broth media (LB) for liquid cultures and on LB-Agar plates (1.5% agar) with respective antibiotics (5 μg/ml chloramphenicol [cm], 100 μg/ml spectinomycin [spec], 25 μg/ml lincomycin and 1 μg/ml erythromycin [MLS], 25 μg/ml kanamycin [kan]) and supplements (0.01 mM, 0.05 mM, 0.1 mM, or 0.5 mM Isopropyl-β-D-thiogalactopyranosid [IPTG]). For liquid cultures of *E. coli*, 100 μg/ml ampicillin [amp] was added to the media and the culture was incubated on a shaking platform with 200 rpm at 37°C. Liquid cultures of *B. subtilis* were grown at 30°C on a shaking platform (200 rpm) supplemented with the required antibiotics. To verify the successful integration at the *amyE*-locus in *B. subtilis*, LB-Agar plates were supplemented with 1% starch (w/v). The expression from the *amyE-* locus with a *hyperspanc* promotor was induced by IPTG. For competence induction in *B. subtilis* the modified competence medium according to Spizizen was used [39]. A volume of 100 ml of 10x MC-media was composed of: 14.01 g K_2_HPO_4_ x 3 H20, 5.24 g KH_2_PO_4_, 20 g Glucose, 10 ml trisodium citrate (300 mM), 1 ml ferric ammonium citrate (22 mg/ml), 1 g casein hydrolysate, 2 g potassium glutamate. Sterile filtered media was stored at −20°C. For inoculation of a 10 ml culture, 1 ml of 10x MC media, 0.333 ml of sterile 1 M MgCl_2_, the respective antibiotics or supplements were added and filled up with autoclaved water. Cells were grown to competence on a shaking platform (200 rpm) at 37°C.

### Strain constructions

To investigate point mutations of ComGC, the gene was amplified from chromosomal DNA of PY79 with primers listed in table S1. The gene was cloned into pDR111 using *NheI* and *SphI* sites, for an ectopic expression at the *amyE*-site (PLASMIDS SEE TABLE S2). The plasmid does not contain a ribosomal binding site (RBS); therefore, 24 bp encoding a RBS were added N-terminally to all constructs by a second vector-PCR (sequence, including **RBS**: GATTAACTAATA**AGGAGG**ACAAAC. The presence of the RBS was verified by sequencing. To generate point mutations in the *comGC* plasmid, DNA was isolated and a vector-PCR was used. Primer-pairs were designed back-to back with a size of 20 bp each, including the mutation at the beginning of the forward primer. Afterwards the DNA was de-phosphorylated, ligated, and in a further step template DNA was removed by *DpnI*. The plasmid was transferred into chemically competent *E. coli* DH5α. Competent PY79 cells were transformed with plasmid DNA and integration at the *amyE-site* was verified by an amylase assay on starch agar. To this end, plates were covered with 5 ml of Iodine-potassium iodide solution (Carl Roth) and incubated for 10 min at room temperature. Each time the assay was performed, a positive and a negative control were included. List of strains see table S3.

To investigate point mutations at the original locus of *comGC*, the vector *pMAD* [40] was used. For generation of the PCR products and the point mutation, Gibson-Assembly and Q5-site directed mutagenesis was used. Primers were designed with the NEB primer design tool (New England Biolabs).

### Transformation of B. subtilis & assays of transformation frequency

Freshly restreaked strains on LB-Agar plates were inoculated as a 3 ml LB-overnight culture, which was grown at 37°C and 200 rpm supplemented with inducing agents and antibiotics. The following day, a culture was inoculated to an OD_600_ of 0.08, and grown at 200 rpm until at 37°C the culture reached an OD_600_ of 1.5 in MC-media (see growth). For transformation 0.5-1.5 ug of chromosomal or plasmid DNA was added to the cells and incubated for 1-2 h at 37°C and 200 rpm. Transformants were obtained after an incubation time at 30 degrees for 48 h.

For transformation frequency assays, 0.5 ml of competent cells were incubated with 0.5 μg of chromosomal DNA, to have a final concentration of 1 μg/ml for 1 h at 37°C and 200 rpm. A serial dilution was made, and 10^-1^, 10^-2^ or 10^-3^ dilutions were plated on LB-Agar plates with respective antibiotics to obtain colony forming units (CFU). A 10^-6^ dilution was used to obtain the number of viable cells. By dividing the CFU/ml by the number of viable cells/ml/μg DNA, efficiency was calculated. For each strain a technical and biological duplicate was performed. Visualisation of data was generated with GraphPad Prism6 (GraphPad Software, San Diego, California, USA). To test the influence of Alexa Fluor 488 C5 Maleimide (AF488-C5 maleimide) on the transformation activity of the strains, cells were treated with the stain before addition of chromosomal DNA.

### Fluorescence staining of DNA

DNA was fluorescently labelled according to a modified protocol of Boonstra *et al*. [41]. A PCR product containing an Erythromycin resistance cassette flanked by homologous *B. subtilis* regions with a size of 2300 bp was employed for staining by using the primers pMB013 and pMB014. 5-[3-aminoallyl]-2’-deoxyuridine-5’-triphosphate (aminoallyl-dUTP, Thermo Scientific™) was incorporated into a PCR by the use of a DreamTaq^TM^ DNA Polymerase (Thermo Scientific™) and template DNA was removed by using *DpnI* for 1.5 h at 37°C. The PCR product was purified and eluted in 50 μl of 0.1 M NaHCO3 solution at a pH 9. DNA was stained by using Dylight562 NHS Ester (Thermo Scientific™) which binds to aminoallyl-dUTP. For reaction with the stain, an aminoallyl-ester with 10x excess of dye was added to the PCR product. In a further step the PCR product was labelled for 3h at 25°C in the dark. The amount of dye was calculated by:

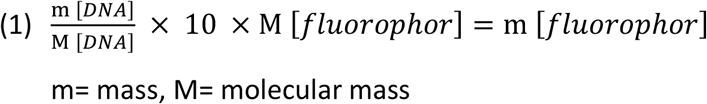

The DNA was purified by using a PCR purification kit (Quiagen) with an additional washing step. Labelling of the PCR product was checked by fluorescence signal with the Biomolecular Imager (Typhoon, Amersham) of a 1% agarose-gel. Labelled PCR-products were stored at −20°C under exclusion of light.

### Staining of pilus structures

Competent cells were centrifuged with an Eppendorf centrifuge 5424 at 8000 rpm for 2 minutes, washed with 1 x PBS (pH 7) and subsequently labelled with 25 μg/ml AF488 C5 Maleimide (Invitrogen) for 10 min in the dark followed by at least two additional washing steps. The pellet was resuspended in supernatant from cells grown to competence.

### Epifluorescence microscopy

Cells grown to competence were incubated at 37°C and 200 rpm to an OD_600_ of 1.5. Cells were transferred onto 1% agarose pads (w/v) containing conditioned medium, generated by sandwiching 100 μl of melted agarose between two coverslips (12 mm, Menzel). 3 μl of the culture were spotted onto a round coverslip (25 mm, Marienfeld). For wide field image acquisition a Zeiss Observer A1 microscope (Carl Zeiss, Oberkochen, Germany) with an oil immersion objective (100× magnification, 1.45 numerical aperture, alpha Plan-FLUAR; Carl Zeiss, Oberkochen, Germany) was used with a charge-coupled-device (CCD) camera (CoolSNAP EZ; Photometrics, AZ, USA) and an HXP 120 metal halide fluorescence illumination. A YFP filter (ET500/20 EX T515lp ET535/30 EM) was used for epifluorescence or 488 nm laser excitation.

Samples were illuminated for 0.5 to 2 s, dependent on signal intensity, at the midcell plane. Final editing of images was done with ImageJ2/ FIJI 1.52 [42].

### Single-molecule tracking (SMT)

Cells grown to competence were prepared with AF488-C5 maleimide (see staining of pilus structures). In case of incubation with chromosomal DNA cells were incubated with 20 μg/ml chromosomal DNA (PY79) for 20 minutes as described [27]. Slides and samples for microscopy were prepared as described in Epifluorescence microscopy. For Single molecule tracking (SMT) an inverted microscope (Nikon Ti Eclipse) was used, with a 514 nm laser diode (100 mW, Omicron Laser) with an A=1.49 objective and an EMCCD camera (Hamamatsu) by using an integrated autofocus on the microscope. ImageJ2/ FIJI 1.52 [43] was used for movie cutting and processing. Movies were cut to reach a single molecule level by removing the initial bleaching steps, until the bleaching curve had a remaining decline of less than 10%. Cell meshes were set with oufti [44] and tracks were detected with UTrack [45]. In a further step tracks were analysed with the SMTracker software 2.1 [46]. Diffusion constants and fraction sizes were calculated by a squared displacement analysis (SQD). The SQD is the cumulative probability function of squared displacements.

### SIM Microscopy

Structured illumination microscopy was used for generation 3D reconstructions of cells. Images were acquired with the ZEISS ELYRA PS.1 (Laser lines 561 nm/ 488nm; ANDOR iXon EMCCD; ZEISS objective: alpha Plan-Apochromat 100x/NA 1.46). Reconstructions were generated with the software ZEN-Black. Final editing and 3D-reconstructions of images was done by ImageJ2/ FIJI 1.52 [42].

## Results

### ComGC can be modified into a cysteine-carrying variant retaining transforming activity

We pursued the strategy of cysteine-labelling to visualize a possible pseudopilus or pilus structure in *B. subtilis* cells grown to competence. We mutated several amino acids of ectopically integrated ComGC to cysteine residues, and integrated these into the *amyE* site on the chromosome, driven by an IPTG-inducible promoter (data not shown). We found a change at position 71 to cysteine to retain transformation activity, even when cells were incubated with a fluorescent maleimide stain to label ComGC (Fig. 1). As expected we were not able to obtain transformants in a *ΔcomGC* deletion strain, which we used as a negative control [47]. In a next step, we integrated the corresponding codon for the 71C amino acid change into the original *comGC* gene, and tested for transformability. The mutation did not lead to a reduction in transformation activity, yielding 3.35 x 10^-5^ +/-1,19×10^-5^ (Fig. 1), indicating that the point mutation does not have an effect on the functionality of ComGC. We were surprised that the Alexa Fluor488 C5 Maleimide (termed “AF488-C5 maleimide from here on) - staining procedure generally increased transformation efficiency of all constructs tested, independent of the presence or absence of the ComGC^CYS^ variant (Fig. 1). While the finding shows that staining did not have a negative effect on the functionality of ComGC, we can not offer a conclusive explanation for the effect, other than a possible additive interaction of the fluorescent dye with DNA.

**Figure 1:**
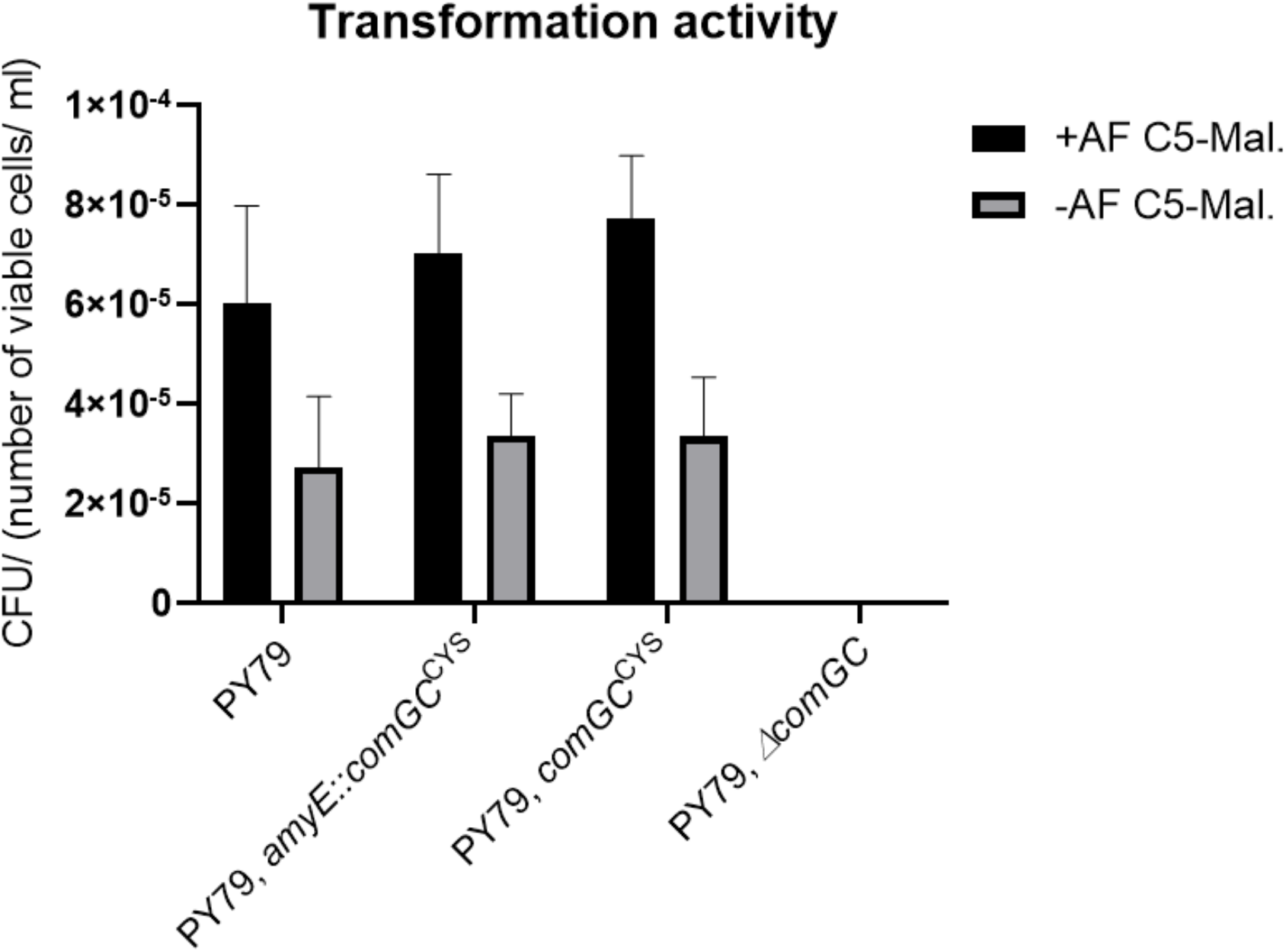
Transformation frequencies of strains encoding ComGC from the *amyE* locus and mutations. The bar plot indicates the CFU (colony forming unit) defined as the number of viable cells per millilitre per microgram added chromosomal DNA. Experiments were done in technical and biological duplicates. Error bars indicate the standard deviations.

Because we anticipated that a mixture of wild type ComGC and AF 488 C5-maleimide-labelled ComGC might be beneficial for pilus formation, based on the idea that too many labels within a pilus might hamper dynamics and thus imaging, we expressed the cysteine variant from the amylase locus, using mild IPTG induction. We found comparable transformation efficiencies like wild type cells (Fig. 1). As a control for the following experiments, we created a strain that ectopically expresses wild type ComGC from the *amyE* locus. For this strain, we detected higher numbers of transformants with an activity of 1.13 x 10^-4^ ± 1.03 x 10^-4^, or 2.16 x 10^-4^ ± 1.84 x 10^-4^ after treatment with AF 488 C5-maleimide (not shown in Fig. 1 because of the high standard deviation). Because of the high standard deviation, we cannot conclusively state that higher amounts of ComGC generally increase efficiency. In spite of the inexplicable effects of increased transformation via staining and ectopic ComGC expression, the experiments suggest that a change of serine at position 71 to cysteine leads to sufficient activity of ComGC to study its *in vivo* localization.

### *Bacillus* cells grown to competence carry a true pilus on their surface

When cells expressing ComGC from an ectopic site were fluorescently labelled by treating the cells with AF488 C5-maleimide, filamentous structures emanating from the cell surface were observed (Fig. 2A), indicating that a pilus-like structure can be visualized in *B. subtilis* cells. Filamentous structures were observed at the cell poles (Fig. 2A, mid panel), at the cell center (Fig. 2A, lower panel) and often more than one structure per cell was visible (Fig. 2B upper panel). The structures were not seen in the negative control, which expresses ComGC from the *amyE* locus of competent cells (Fig. 2B). Instead, only staining of the membrane and of the septum was detectable. Counting of 385 of such cells revealed that none of them formed pilus structures.

**Figure 2.**
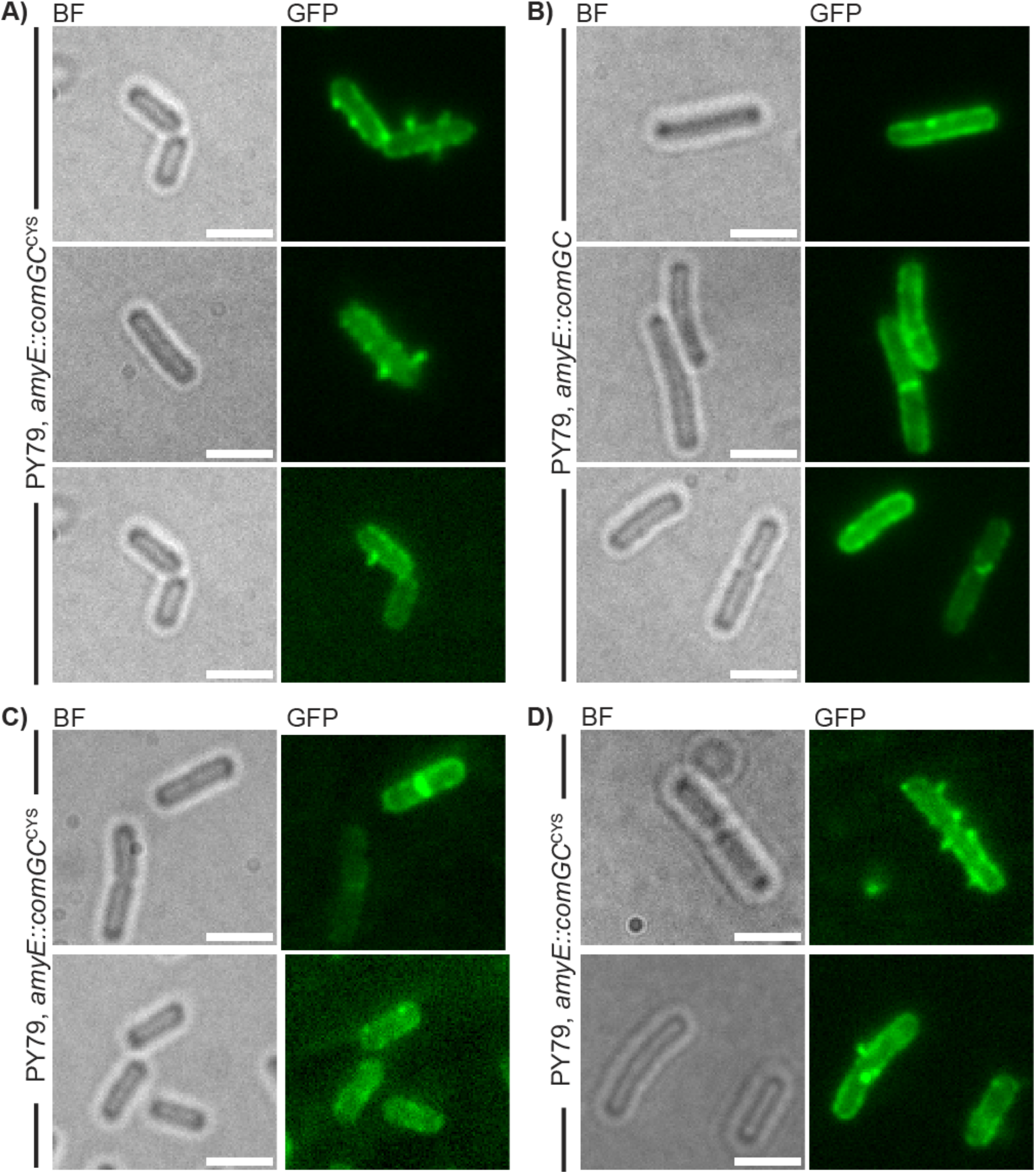
Epifluorescence microscopy of AF488 C5-maleimide stained cells of mutated (PY79 *amyE::comGC^CYS^*) and cells (PY79 *amyE::comGC*) encoding ComGC at the *amyE* locus. With 0.5 mM IPTG induction (panel A and B), without IPTG induction (panel C) and with low (0.01 mM) concentrations of IPTG (panel D). Left images of a panel show bright field images (BF), left panels show epifluorescence pictures (GFP). Scale bars represent 2 μm.

Contrarily, in cells expressing ComG-CYS, 64 cells showed pilus structures out of 318 analyzed cells, which is in accordance with the literature value of 10-20% of cells inducing the state of competence [19]. The formation of the structures was dependent on IPTG, cells incubated without any IPTG did not show any pili (Fig. 2C). To ensure that the formation of the pilus structures is not based on overexpression, we used a low concentration of IPTG (0.01 mM). Even with low induction we were able to see pili, which supports the assumption that the formation was caused by incorporation of ComGC-CYS into existing ComGC-based pili. We wanted to know if pilus formation is dependent on external DNA. For this, we incubated a ComGC-CYS AF488 C5-maleimide stained strain with chromosomal DNA and quantified the number of pili in these cells by using Epifluorescence microscopy. By counting ~200 cells we were not able to observe any striking change in pilus quantity by treatment with chromosomal DNA, which lead to a slight decrease of 1.2 fold in formation of pilus structures, suggesting that pilus formation is not depending on the availability of DNA in a culture.

We used super-resolution Structured Illumination Microscopy (SIM), reaching a resolution of 125 x 125 nm in x and y direction, to generate 3D reconstructions of cells. With this technology, it is feasible to determine if a cell envelope-spanning machinery is exposed from the surface of *B. subtilis*. Z-stacks of stained cells were obtained, and signal was seen at sites emanating away from the cell surface (Fig 3, supplementary movie S1). Supplementary movie S2 displays Z-stacks of AF488 C5-maleimide stained cells. Staining of cells expressing ComGC^CYS^ from the original gene locus did not show structures that could be well-resolved (data not shown). We favour the view that pili can be visualized best when a mixture of native (non-labelled) and cysteine-mutant (labelled) ComGC is expressed in the cells. We further used SIM to measure the length of pili, which was 505 ± 155 nm (n = 35) under full induction (0.5 mM) of the hyperspank promoter. To ensure, that length is not caused by overexpression of labelled ComGC, we also measured pilus length using low (0.01 mM) IPTG induction. Pilus length was 539 +/− 40 nm (n = 40), showing that length does not correlate with induction levels. Thus, keeping in mind the caveat that we used a merodiploid strain to measure average pilus length, we believe it is safe to argue that *B. subtilis* pili are considerably shorter than those observed for gram-negative bacteria [32, 33]. These findings support the idea that DNA uptake through the cell wall in *B. subtilis* is not restricted to the cell poles, but occurs all over the cell surface.

**Figure 3.**
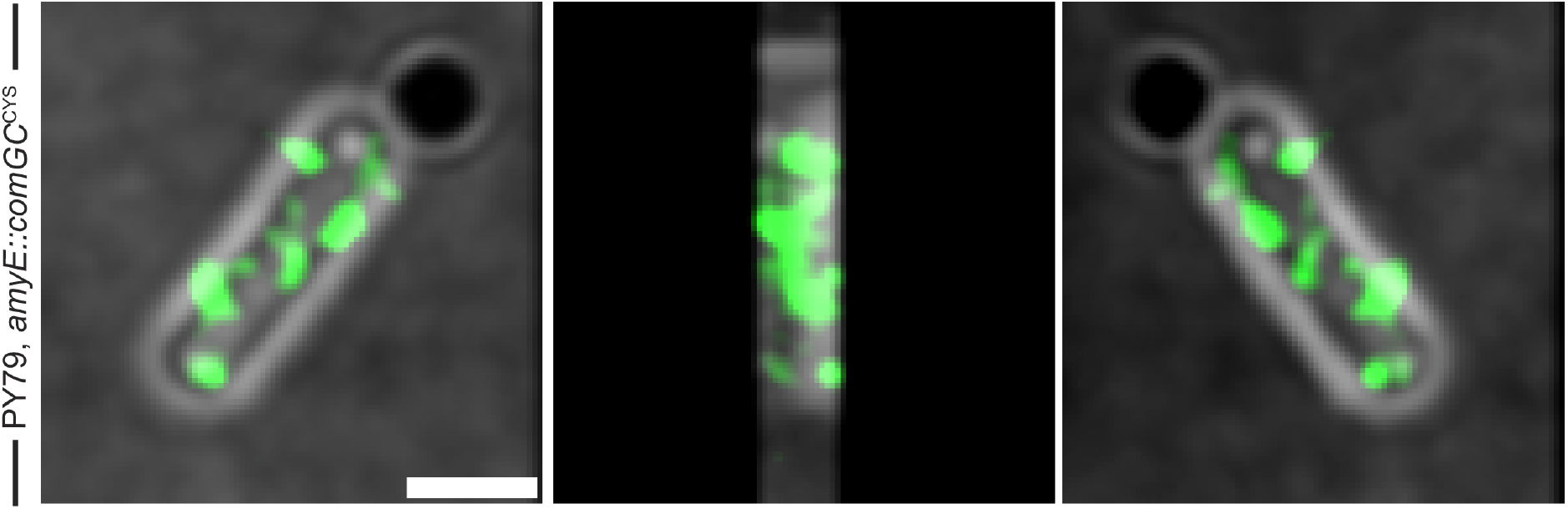
3D-reconstructions of Structured illumination microscopy of AF488 C5-maleimide-stained cells encoding ComGC^CYS^ from the *amyE-locus*. A) Left panel shows 0°C, middle panel 180° and right panel 270°. Corresponding movie S1. Scale bars represent 2 μm.

### Pili are dynamic structures that interact with external DNA

Competence pili as well as pili used for surface-based motility are known for dynamic extension/retraction periods, in which pili grow by addition of pilins to the base of the pilus, or by detachment of pilin subunits from the base, and their presumed diffusion back into the cell membrane. We tested if *B. subtilis* pili show such time-dependent remodelling, using 20 seconds time lapse experiments. Fig. 4A shows an example of a cell displaying multiple pili; the orange triangle indicated a pilus that is retracting (movie S3). Fig. 4B highlights a cell, in which a pilus that is extending (yellow triangle), and a pilus indicated by the orange triangle retracts within the last two time intervals (see movie S4). Of note, we did not observe considerable lateral motion of pili along the cell surface. These data suggest that pili are stably anchored, either at the cell membrane or, more likely, in the cell wall, and are predominantly static on a scale of 3 to 4 minutes, while a significant number displays dynamics in a second-time scale, expected for a structure that pulls external DNA through the cell wall.

**Figure 4.**
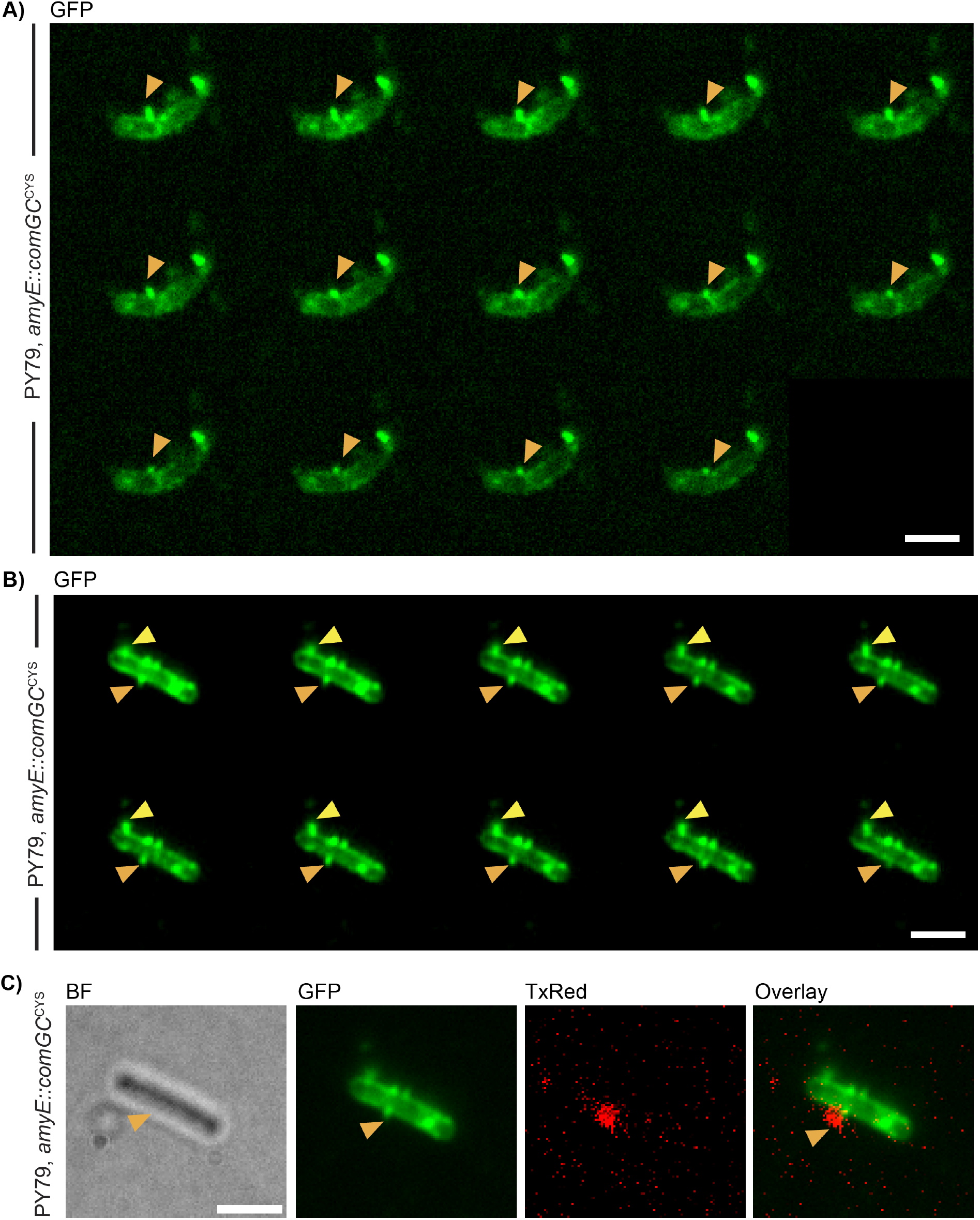
Dynamics of AF488 C5-maleimide-stained competence pili. **A+B)** Images of time-laps microscopy were acquired every 20 s. Yellow arrow indicates pilus formation, orange pilus retraction. Movies were acquired in the GFP-channel. **C)** Colocalization of fluorescently labelled DNA with C5 Maleimide-stained competence pili. Scale bars 2 μm.

To test if an interaction between DNA added to the culture and competence pili can be observed, we used fluorescently labelled DNA in strains expressing ComGC^CYS^. Labelled DNA was checked for fluorescence signal with a biomolecular imager, which showed fluorescence signal at 2300 bp corresponding to the size of the amplified PCR product, including an erythromycin resistance cassette and homologous regions of the *B. subtilis* chromosome [28]. Fluorescently labelled DNA was incubated with AF488 C5-maleimide-stained cells that were treated with DNase in a further step to ensure that no additional, exogenous DNA was present in the culture. Indeed, we were able to detect fluorescence signal for the labelled DNA that colocalized with pilus structures (Fig. 4C). because this event was seen in few cells, we would assume that a colocalization of taken up DNA by a pilus structure is a rare event. Of note, fluorescently labeled DNA yielded a normal number of transformants, comparable to non-labelled DNA [28, 41].

### ComGC-CYS is diffusing along the membrane in SMT

Pilin subunits of pili are expected to diffuse through the cell membrane, and to assemble into a polymeric structure. Because we are not aware that dynamics of pilin have been visualized at a single molecule level, we set out to obtain more information on the dynamics of competence pili and performed single molecule tracking experiments with ComGC-CYS, expressed at very low levels from the *amyE* site. Because transformation efficiency was high in this strain, and because it is reasonable to assume that ComGC-CYS mixes with wild type ComGC and forms joint pili, we tracked ComGC under these conditions, which appear to most closely resemble native conditions. In other words, because ComGC-CYS is visibly incorporated into pilus structures, it should serve as good proxy to ComGC dynamics *in vivo*. SMT allows to follow the motion of freely diffusive molecules in living bacteria with a resolution of 40 nm and less. For tracking of single molecules, we used a 30 ms stream acquisition for 2500 frames. SMTracker 2.0 was used for data analysis of single molecule tracks that were generated with U-track [45]. Cell meshes for analysis with SMTracker were set with the software oufti [44]. Continuous tracks of at least five steps were used for data analysis.

At first, we localised ComGC-CYS molecule by projection of tracks into a normalized (medium-size) *B. subtilis* cell with an average size of 3 x 1 μm. Red tracks in Fig 5A represent molecules moving in confined motion, within an area of 108.7 nm in diameter, in cells grown to competence. The confined area corresponds to 3 times the localization error determined from mean squared displacement (MSD) analyses. Confined motion was restricted to the cell membrane, and the septum at mid-cell, as expected for a membrane protein (Fig. 5B). Confined tracks likely represent ComGC-CYS molecules engaged in pilus formation. Freely diffusing molecules are shown in blue; note that the cell is over-saturated with these tracks in order to obtain a sufficient number of confined tracks. Green tracks represent molecules showing transitions between diffusive and confined motion, and largely overlap with areas of confined motion.

**Figure 5.**
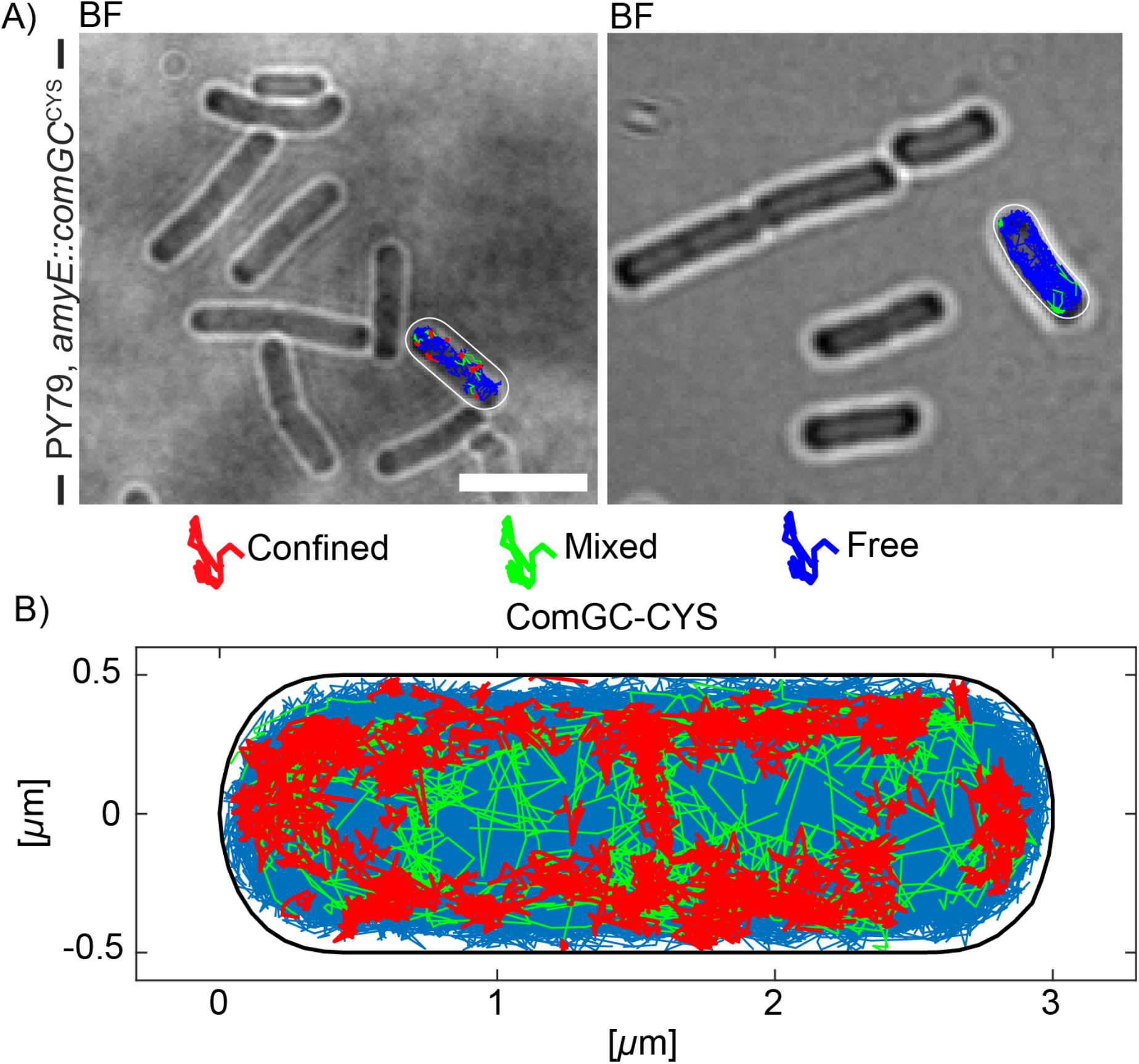
Localization of Single molecule tracks of ComGC-CYS treated with AF488-C5 maleimide. A) Overlay of Confinement map of tracks in representative cells cell and B) Confinement map of confined tracks projected into a standardized *B. subtilis* cell. Confinement radius is 108.7 nm, which corresponds to 3 times the localization error. Confined tracks are indicated in red, green tracks show tracks from a confined to a more mobile movement and *vice versa*, while blue tracks are defined as mobile, moving out of the confinement radius. Scale bar represents 2 μm.

Jump distance (JD) analysis was used to obtain more detailed information about the diffusion constants and fraction sizes of ComGC-CYS. JD analysis describes the Eucledian distance between consecutive detections [48]. A nonlinear regression fit was used to describe the data (Rayleigh fit). By using one Rayleigh fit, the generated data were not well represented (Fig. 6A, upper panel). Two Rayleigh fits had to be used to describe the data sufficiently well (Fig 6B, upper panel). Middle panels show the difference between fitted data and modelled data based on Brownian diffusion. Please note the different Y-axis dimensions, showing that differences were much smaller when using two Rayleigh fits (Fig. 6B). Lower panels present overall fitting of data with modelled data, which shows that two fits leave very little deviation (Fig. 6B), compared with the single fit (Fig. 6A). These analyses suggest that two populations with distinct diffusion constants exist for ComGC-CYS *in vivo*. Using JD data, diffusion constants and fraction sizes of the two populations were determined. Figure 7B visualizes the data shown in Fig. 6B: the size of the bubbles corresponds to the relative size of the population and the height along the Y-axis to the diffusion constant of the protein. The slow population (“static fraction”, Pop1) had a size of around 32% of ComGC-CYS molecules, with a diffusion constant (D1) of 0.042 ± 0.001 μm^2^/s, while the faster and larger population (Pop2) with 68% of ComGC-CYS had a diffusion constant (D2) of 0.51 ± 0.001 μm^2^/s (table 1). The latter is well within the range of diffusion constants obtained for other freely diffusive membrane proteins in *B. subtilis* [49], the prior with a rather static structure. Treatment of PY79 *amyE::comGC* cells with Alexa-Fluor C5-maleimide showed unspecific binding of the stain (Fig. 2B). To control for ComGC-CYS tracks due to unspecific binding of the stain, we performed single-molecule tracking of PY79 *amyE::comGC* with and without staining. Fig. S1 shows than less than 10% tracks were monitored in cells lacking staining and/or the cysteine exchange; non-specific tracks were also considerably more slow than those observed for stained ComGC-CYS (Fig. S1, bubble blots). These data suggest that about a third of ComGC molecules are statically engaged in pili, while two third are diffusive within the membrane, able to be recruited into pilus structures.

**Figure 6.**
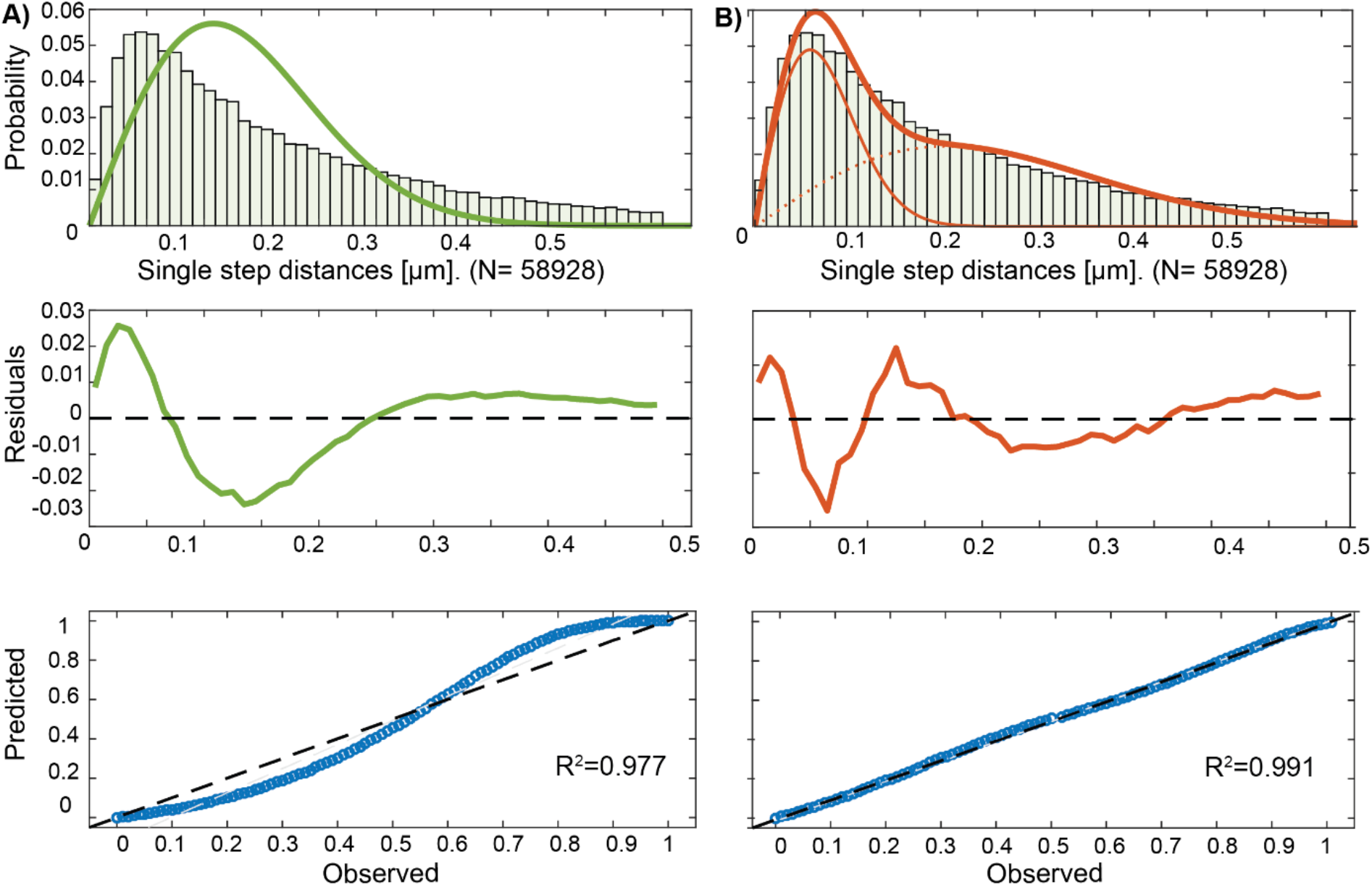
Squared Displacemend analysis (SQD) of ComGC-CYS cells treated with AF488 C5-maleimide in *B. subtilis* cells grown to competence. A and B (upper panel) show Rayleigh fits of the generated data of one and two populations. The plot shows the frequency of diffusion constants depending on the diffusion coefficient of the tracks in a histogram. The probability R^2^ is annotated for each histogram, a two population fit is shown in B, which better describes the data as in A. Lower Panels of A and B show quantile–quantile plots for one (A) and two populations (B) which illustrate how well measured (dashed line) and the modelled (blue line) data fit together.

**Figure 7.**
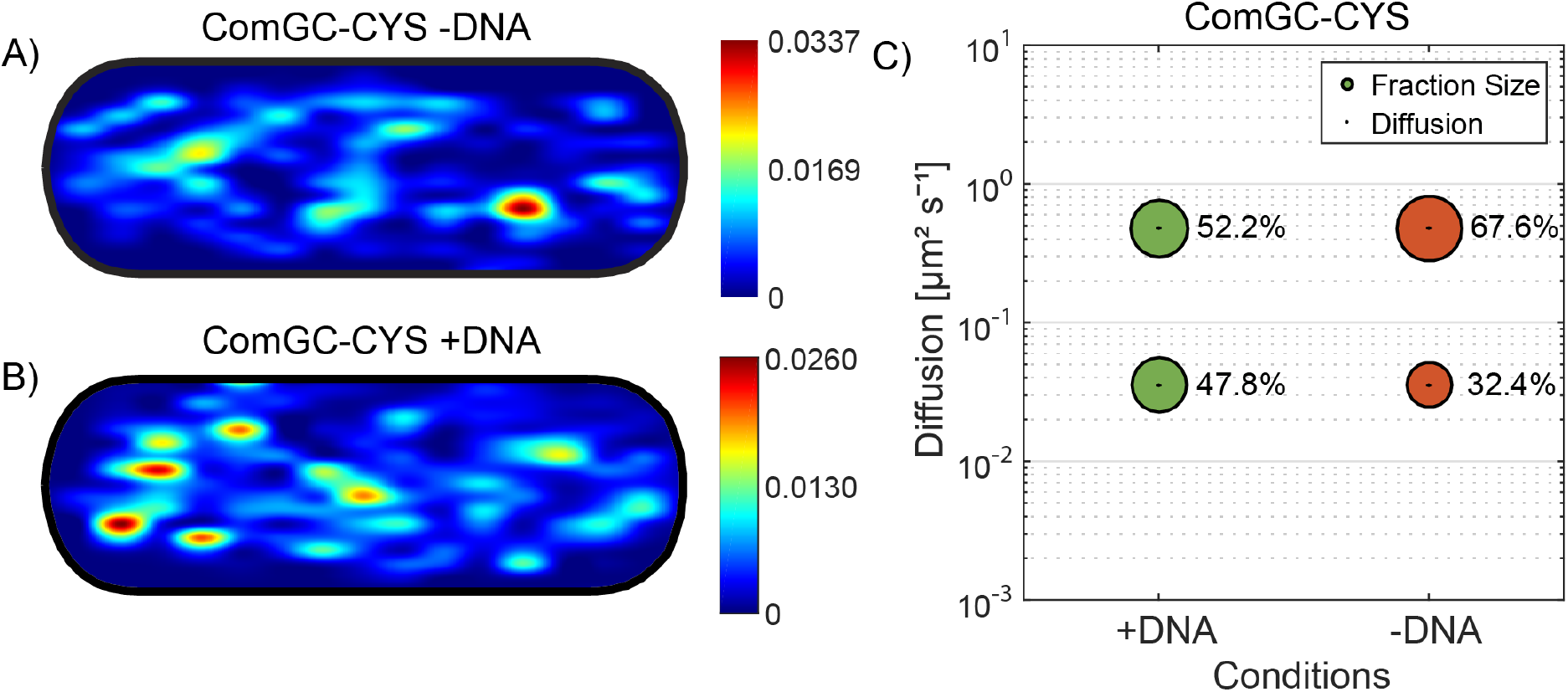
Response of pilus dynamics after addition of DNA. Panel A) shows heat maps of confined tracks of ComGC-CYS stained cells with AF488 C5-maleimide upon addition of DNA (indicated above) in a standardized *B. subtilis cell*. Confinement radius was 108.7 nm. Colour code on the right indicates intensity of signal in presence of protein. B) Bubble plot shows diffusion constants [μm^2^/s] the fraction sizes of populations by setting a simultaneous diffusion constant which is shown in correspondence. Errors correspond to the 95% confidence intervals which are given by “confint” matlab function by using its values that result from the fit. +DNA and -DNA indicates cells which were treated with DNA for 10 minutes.

**Table 1.** Single molecule tracking of ComGC-CYS with Alexa-Fluor C5-maleimide treatment, with/ and without addition of DNA with an incubation time of 10 minutes.

|  | PY79<br>*amyE::comGC<sup>CYS</sup>* - DNA | PY79<br>*amyE::comGC<sup>CYS</sup>* + DNA |
| --- | --- | --- |
| Pop<sub>1</sub> [%] | 32.4 | 47.3 ± 0.005 |
| Pop<sub>2</sub> [%] | 67.6 | 52.7 ± 0.005 |
| D<sub>1</sub> [μm<sup>2</sup>/s] | 0.042 ± 0.001 | 0.038 ± 0 |
| D<sub>2</sub> [μm<sup>2</sup>/s] | 0.51 ± 0.001 | 0.22 ± 0.003 |

### The static fraction of ComGC-CYS increases after addition of chromosomal DNA

We wished to addressed the question of competence pili that interact with DNA show altered dynamics. For this we tested different incubation times of ComGC-CYS with chromosomal DNA. Tracking of ComGC-CYS after 20 minutes incubation time with DNA yielded similar sizes of populations and of the diffusion constants [Fig. S2]. Considering that possibly, most of the added DNA is already taken up in 20 minutes and located either in the periplasm or in the cytosol, we used a decreased incubation time of 10 minutes for following experiments. Also, we ensured that only cells with visible pilus structures were analysed. For this, an epifluorescence image was taken before tracking and only cells with visible exposed surface structures were used for further analysis. Here, the static fraction of ComGC-CYS showed a significant shift of its size from 32.4% to 47.8% (Fig. 7C). Concomitantly, the diffusion constant of the static fraction somewhat decreased from 0.042 to 0.033 ± 0.001 μm^2^/s with addition of DNA, while that of the fast population only mildly decreased from 0.51 to 0.45 ± 0.001 μm^2^/s (table 1).

Moreover, by projecting approximately 500 tracks into a normalized *B. subtilis* cell of 3 x 1 μm size and sorting the tracks which stayed within a confined radius of 108.7 nm we observed a change in the localisation pattern (Fig. 7A). The degree of confined tracks at the cell periphery was visibly increased (note that *B. subtilis* cells are 0.7 μm wide), likely representing ComGC within active pili (Fig. 7B), in agreement with the 47% increase in the static population (Fig. 7C). However, this change in confinement is not reflected in the number of pili visible by SIM or epifluorescence microscopy, which was only 1.2 fold higher in cells after addition of DNA. We interpret these findings to indicate stochastic, DNA-independent polymerization of competence pili, and an increased time ComGC spends in its polymerized form, suggesting a longer time for retraction due to bound DNA.

## Discussion

*B. subtilis* cells take up DNA into the cytosol at a single, and less often at both cell poles [26]. This localized transport through the ComEC channel leads to the assembly of recombination proteins at a single subcellular cite, possibly facilitating search for homology on the chromosome of incoming DNA but a two-dimensional process [50]. It has been unclear of DNA is also taken up from the environment at a single polar site, which would decrease the probability of interaction with DNA, and if uptake occurs via a pseudopilus, i.e. a short structure that opens a gate through the wall, or via a true pilus, such as in *S. pneumoniae* cells, which can “fish” DNA along a longer distance away from cells [33].

It has long been known that *B. subtilis* cells contain an operon of pilin-like genes that plays an essential role in DNA uptake. The encoded ComG proteins have similarities to proteins forming Type IV-pili and type II-secretion systems of gram-positive and gram-negative bacteria [51]. Degradation of the cell wall relieves the essentiality of the operon, and biochemical experiments have shown that the major pilin, ComGC, forms a complex of a size of at least 400 to 1000 kDa, in conjunction with minor pilins [29, 51]. It was also shown that these proteins might facilitate the interaction of DNA with the DNA receptor [51].

Fusions of mCherry with ComGC showed no functionality in DNA uptake. By using Immunofluorescence microscopy and a FlaSH-Tag, ComGC was shown to localize to the cell poles and the cell center [52]. Therefore, it was proposed that cells may contain just one active, assembled competence machinery and freely diffusive single competence proteins [52]. By using a pilus labelling method we show that indeed, *B. subtilis* cells possess a competence pilus, a machinery spanning through the cell envelope, which was detected in 20% of cells grown to competence. The length of the structures was analysed by structured illumination microscopy, a super-resolution method by which a resolution of down to 125 nm can be reached. Structures with an average of 500 nm length were measured, indicating a much shorter structure then reported for gram-negative bacteria like *V. cholerae* [32].

Intriguingly, the competence machinery has been shown to be located at one or both cell poles, suggesting that DNA uptake occurs at specific sites in competent cells [26]. Our results reveal that the pilus in *B. subtilis* is located at the cell poles, like it was reported, but also at any other site within the cell envelope, indicating that DNA uptake through the cell wall is not limited to the poles, in contrast to intracellular uptake through the cell membrane (Fig. 8). Also, the presence of more than one competence pilus revealed that there can be more than one working structure for DNA uptake. This was already proposed by Boonstra et al. [41]: the authors showed that that fluorescently labelled DNA also was taken up at the cell center, and not just at the poles. Uptake at any site on the cell surface, using several pili, increases the efficiency of DNA uptake from the environment, while transport into the cytosol at a single, polar site, may organize RecA filaments that guide incoming DNA to the corresponding sites in the genome [50] (Fig. 8). Likewise, storage of incoming DNA within the periplasm may increase transformation efficiency, and allow for slow transport of ssDNA into the cytosol during the concomitant possibility to rapid uptake of dsDNA through the cell wall.

**Figure 8.**
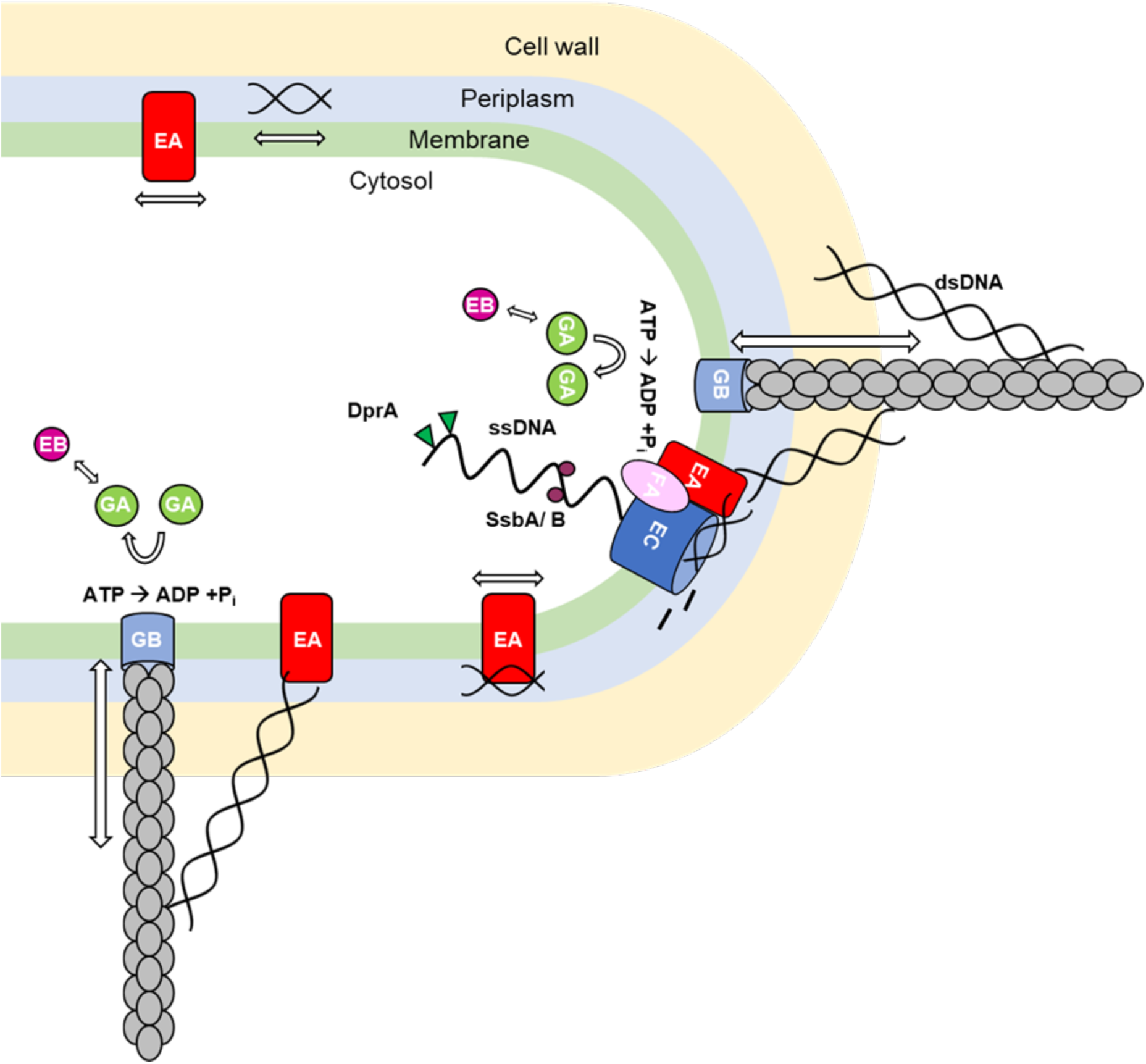
Model of DNA uptake in *B. subtilis* as a two-tier process. The periplasmatic uptake of double stranded (dsDNA) occurs over the whole cell surface via a true pilus structure, showing dynamics assembly and disassembly. The anchor of the structure might be ComGB (light blue). Via pilus retraction, bound DNA will be pulled into the periplasm where it will bind to the DNA receptor ComEA (red), and diffuse throughout the periplasm. Once reaching the poles, it can be taken up by ComEC (dark blue). ComFA (violet) might provide the energy for the uptake. Direct passage through the pilus, via ComEA to ComEC, may be possible at the pole. ComEC processes double stranded (ds) DNA into single stranded (ss) DNA, which once inside the cytosol will be coated by single strand DNA binding proteins SsbA, SsbB (purple) and DprA (green). This is followed by an integration into the chromosome via dynamic RecA filaments.

We also studied the dynamics of pilin ComGC at a single molecule level, for which we have not found earlier data in the literature. We found that as expected from models on pilus formation, a larger pool of ComGC freely diffuses within the cell membrane, while a third of the molecules is statically positioned at the membrane, likely bound within the pilus polymer (Fig. 9). Incubation with chromosomal DNA resulted in a change in the dynamics of ComGC, by an increase of the population size for the static fraction. This suggests that either, more pili are formed in response to DNA uptake, in agreement with our finding of more static structures formed in cells after addition of DNA, for which we have not found any evidence, or that ComEC stays longer within a pilus structure that is pulling DNA across the cell wall, or both. We found an increase in the static ComEC population 10 minutes after addition of DNA, but in contrast, a longer incubation time of 20 minutes for DNA uptake did not lead to a significant change in fraction sizes and diffusion constants. We interpret these finding to suggest that probably, most of the externally added DNA was already taken up within 20 minutes, indicating that the process happens efficiently within a relatively short time period.

**Figure 9.**
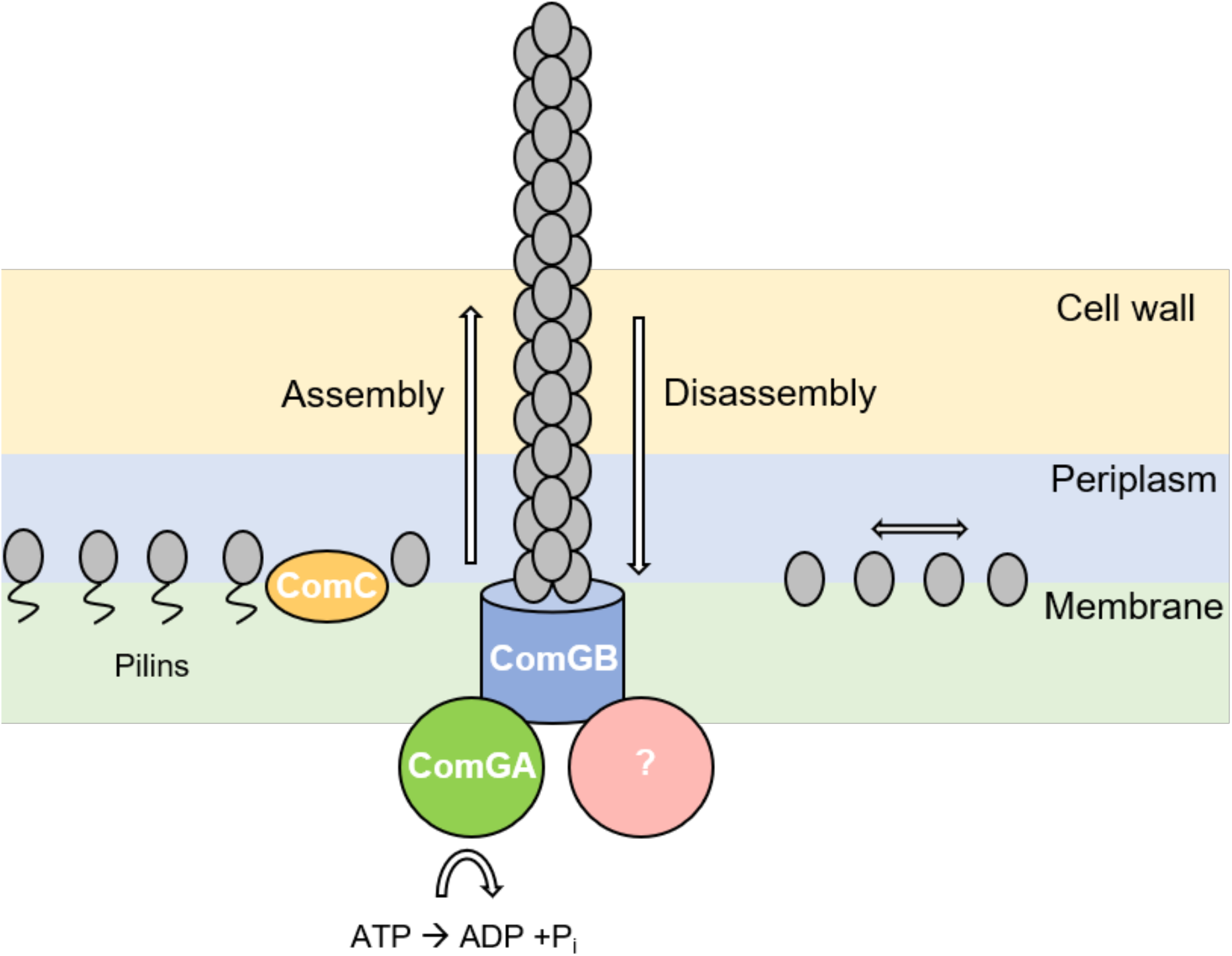
Model for extension and retraction of the competence pilus in *B. subtilis*. Pre-pilins are processed by the prepilin peptidase ComC (yellow) and are assembled into a competence pilus, for which ComGB (blue) might be the platform protein. The energy could be provided by assembly-ATPase ComGA (green), while a retraction factor, or retraction activity, is yet to be identified (purple). Processed ComGC (grey) diffuses within the membrane, while some of the molecules, after having assembled into a competence pilus, are positioned bound within the pilus polymer, represented the static single molecule fraction.

Single molecule experiments have shown that *B. subtilis* cells pull on external DNA with a force characteristic of that typically reported for type IV-pilus retraction [53]. Retraction of pili containing labelled ComGC in our study occurred within seconds to few minutes, in good agreement with single molecule experiments. How might the pilus bind to dsDNA? Purified pilins from *Thermus thermophilus* and from *Streptococcus pneumoniae* have been shown to bind non-specifically to dsDNA [54, 55], possibly forming a tip-structure [56], while pilis from *Neisseria meningitidis* even binds with higher affinity to DNA-uptake sequence (DUS) motif abundant in this species genome [57, 58]. Non-specific DNA binding to *B. subtilis* cells is reduced in *comGC* mutant cells [59], suggesting that the competence pilus also has affinity to dsDNA. We have not been able to find binding of purified, individual pilins to DNA (unpublished results), so it remains to be elucidated how DNA is attached to the pilus for its passage through the cell wall.

Taken together, our data are in favour of a model in which the DNA uptake is a two-tier process in which uptake into the periplasm, in which DNA can freely diffuse bound to ComEA, occurs over the whole cell surface, while cytosolic uptake is limited to the poles, where binding of DNA recombination proteins guides incoming ssDNA onto the nucleoid(s) for homology search.

## Acknowledgements

This work was supported by the Deutsche Forschungsgemeinschaft.

## Supporting Information captions

**Movie S1**

3D reconstruction of an AF488-C5 maleimide stained *B. subtilis* cell grown to competence expressing ComGC^CYS^ from an ectopic site on the chromosome. Movie played with a frame rate of 10 frames per seconds (fps).

**Movie S2**

Reconstruction of a Z-stacks of an AF488-C5 maleimide stained *B. subtilis* cell grown to competence expressing ComGC^CYS^. GIF is shown with 5 frames/s.

**Movie S3**

Time lapse (20 second intervals) of pilus structures of AF488-C5 maleimide stained *B. subtilis* cells grown to competence expressing ComGC^CYS^.

**Movie S4**

Retraction and Extension of pilus structure of AF488-C5 maleimide stained *B. subtilis* cell. Time lapse (20 second intervals) of pilus structures of AF488-C5 maleimide stained *B. subtilis* cells grown to competence expressing ComGC^CYS^.

**Figure S1** Single molecule tracking of ComGC and ComGC-CYS strains with and without treatment with AF488-C5 maleimide.

**Figure S2** Single molecule tracking of ComGC-CYS with and without incubation of DNA.

**Table S1** Oligonucleotides used in this study

**Table S2** Plasmids used in this study.

**Table S3** *E. coli* and *B. subtilis* strains used in this study

**Table S4** Single molecule tracking of ComGC-CYS with/ and without addition of DNA with an incubation time of 20 minutes

## Notes

### Competing Interest Statement

The authors have declared no competing interest.

